# Temporal kinetics of brain state effects on visual perception

**DOI:** 10.1101/2024.08.02.606289

**Authors:** Paul Schmid, Timon Klein, Piotr Minakowski, Sebastian Sager, Christoph Reichert, Robert T. Knight, Stefan Dürschmid

## Abstract

We investigated the effects of brain states on human perception and early visual response comparing focused wakefulness (ON state) to external inattention (OFF state). In two experiments, we investigated the temporal kinetics of brain states changes during stimulus processing and assessed fluctuations across extended periods of time. We used a classifier to distinguish between these states on a single trial level using theta activity in MEG sensors. We found that participants shifted from an ON to an OFF state as rapidly as two seconds. Visual target discrimination was comparable in both states, but reaction times were slower and more variable during the OFF state. Broad band high-frequency activity (BHA) recorded in MEG sensors covering the occipital cortex tracked target grating orientation. BHA was reduced during the OFF state but participants were still able to distinguish sensory information highlighting the role of BHA in visual perception across cognitive brain states.

---

Brain state effects on perceptual responses to sensory input are most evident in the decreased responsiveness observed during sleep or anesthesia ^1^. During wakefulness, humans undergo rapid state changes, frequently transitioning between focused wakefulness (ON state) and inattention to external events (OFF state) ^2^. OFF-state refers to a focus on internal thoughts ^3^ and includes the concept of mindwandering (MW). MW is associated with an increase in perceptual errors ^4,5^ and diminished scalp EEG responses ^6–8^. These findings are interpreted to be due to reduced sensory representation ^9^. However, the neural basis of sensory representation during the OFF state is unclear since differences in the perceptibility of physical changes to visual inputs were not assessed. Furthermore, amplitude reduction in low frequency event-related potentials (lfERPs) typically used in brain state research ^6–8^ does not differentiate between subthreshold vs. suprathreshold cortical sensory representation limiting understanding of the neural basis during transitions between ON and OFF states.

In contrast to lfERPs, broadband high-frequency activity (BHA; 80-150 Hz), is a prominent signal in human intracranial recordings reflecting local neural activity ^10^. BHA visual responses are proposed to index sensory evidence (feedforward) and perceptual awareness (feedback) ^11–14^. Previous studies show that BHA modulation supports decoding of perceptual content independent of the BHA amplitude strength^15,16^. We performed two experiments examining brain state effects on visual perception and early visual BHA. In the first experiment we used a target grating orientation discrimination task. We combined magnetoencephalographic recordings (MEG) and machine learning methods to decode brain states—specifically, ON or OFF-states—at the single-trial level. We trained a classifier using subjectively determined levels of ON and OFF states to predict unlabeled brain states revealing rapid transitions from ON to OFF within seconds. Behaviorally we found no differences in grating orientation discrimination between the ON and OFF states. However, subjects exhibited longer latency and more variable reaction times during the OFF state in accord with prior scalp EEG/behavioral studies. In the second experiment, we compared the temporal modulation of BHA during brain states with the initial feedforward sweep to V1 measured in the C1 evoked response (<100 msec). Contrary to the expected overlap in latency between BHA and C1, the BHA, peaked later and was more sustained than the C1 supporting a role in sensory feedback. Across both experiments BHA was reduced during the OFF state but represented sensory evidence manifested by intact target orientation detection independent of BHA amplitude.

## Results

Combining MEG recordings and machine learning methods in 20 participants in experiment 1, we investigated the relationship between BHA and variations in target orientation and whether this association remained consistent across different brain states. Participants were presented with a visual search array composed of red, green and blue tilted grating patterns displayed on a gray background (see ***Figure 1A***). Red and green gratings alternated as targets and distractors randomly between blocks, while blue gratings served as nontargets. The task was to report via button press whether the target was tilted to the left or the right side. Thought probes were presented pseudorandomly in 20% of the trials and participants rated their level of attention to the trial immediately preceding the probe. Initially, we performed a single-trial classification to identify the frequency band that most accurately reflected the individual ratio of ON and OFF states to test transition times between ON and OFF. We further analyzed whether participants’ behavioral performance and BHA response varied as a function of brain states (see ***Results I – VII***). The results sections correspond to the enumerated methods sections across both experiments (see ***Methods*** for details).

**Figure 1:**
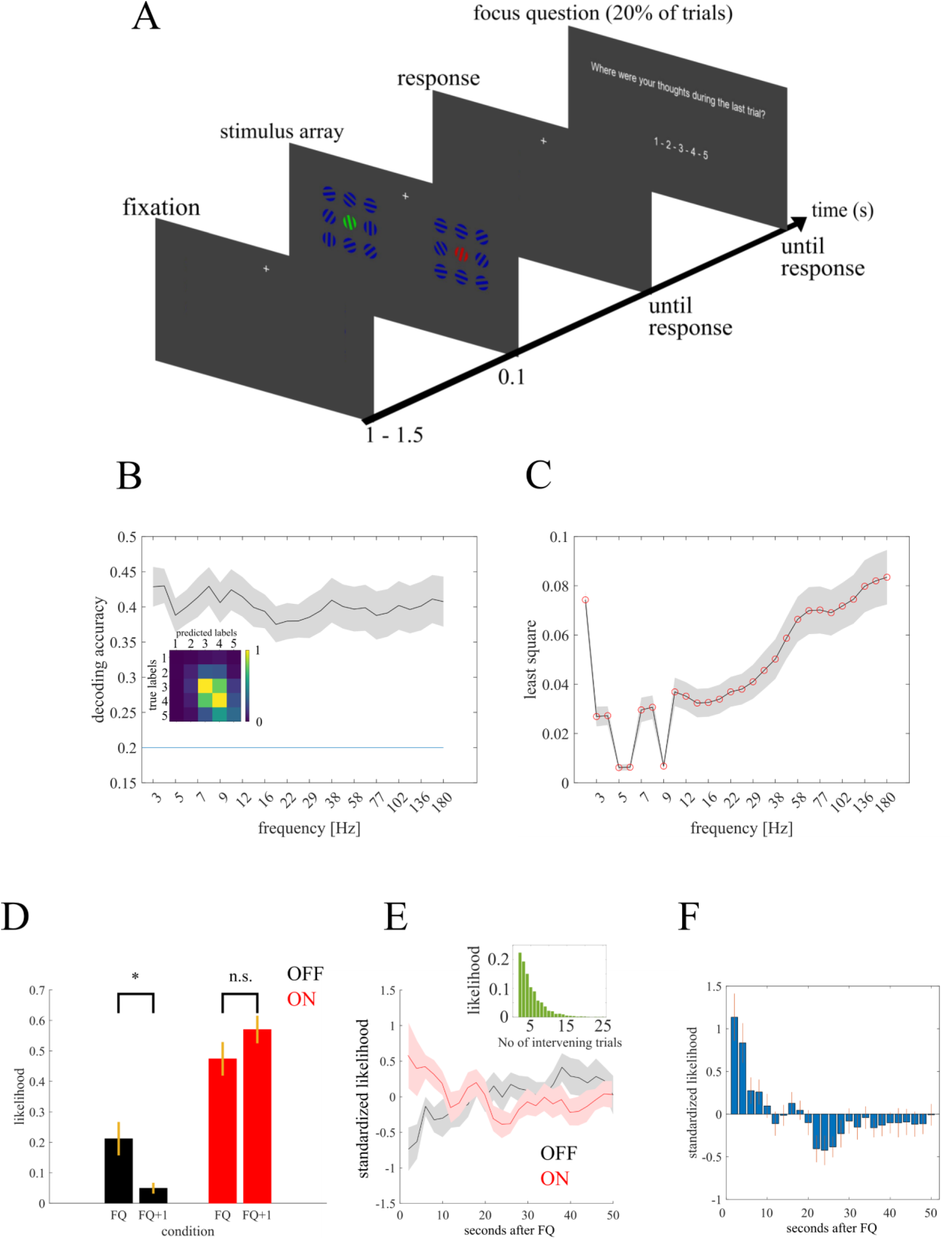
Paradigm and Decoding results. ***A*** Depiction of the paradigm. Single trial with focus question. Participants were instructed to attend either only to the red or green grating - and ignore all other items - and report via button press towards which side the target was tilted. Target gratings could be tilted left or right in ten steps of 1.5°, with the smallest tilt being 1.5° and the maximal tilt being 15° from the vertical axis. Orientation and tilt angle of the non-target and distractor gratings varied randomly***. B*** shows the decoding accuracy (DA) for each frequency band. The shaded area gives the standard error across subjects. ***C*** shows how well the different frequency bands reflect individual OFF-state ratios. The smaller the least square criterion, the better the prediction. Frequencies with center frequencies between 5 and 6 Hz (θ) and 9 Hz (α) best reflect the individual tendency to MW. ***D*** Following thought probes, OFF ratings decrease but ON ratings remain on a stable level indicating that MW is reduced after the thought probes and subjects are focused on the task in the first trial after the focus query. ***E*** while the likelihood of ON labels (red curve) decreases over trials (50 sec with a resolution of 2 sec), the likelihood of OFF labels increases. Shaded areas denote the standard error across subjects. The small inset shows the likelihood distribution of the number of trials between two consecutive thought probes. ***F*** the estimate of ON likelihood measure (ON-label and sign reversed OFF-label likelihood) shows a rapid decline over trials and a significant drop is seen from the first to the second trial after the focus query indicating a drop within 2 seconds.

### I-Representation of individual ON vs. OFF ratio

The likelihood of the five different categories differed (*F*_4.85_ = 4.74; *P* = .0017) with more ON (4: 30.92%; 5: 16.47%) than MID (3: 31.41%) and OFF ratings (1: 9.99%; 2: 11.21%). We tested which frequency band had a similar distribution for predicted labels. We compared the distribution of ON-MID-OFF, resulting from the responses of the participants, with the predicted distributions of each frequency band. The least square criterion provides a parameter indicating how similar the distributions are. Low values represent high similarity. Both θ (center frequencies 5 – 6 Hz) and α (center frequency 9 Hz) showed the strongest similarity with the observed distribution reflecting the best interindividual differences in MW ratings. Given θ activity showed better decoding accuracy (see ***Figure 1B&C***), the following analysis steps are based on the decoding of brain states using theta activity.

### II-Effectiveness of thought probes

We examined ON and OFF ratings between FQ and FQ+1 trials and observed a main effect for factor trial type (*F*_1,19_ = 72.6, *P* < .0001; see ***Figure 1D***) and an interaction effect (*F*_1,19_ = 7.9; *P* < .0001). Post-hoc tests revealed a significant decrease of OFF ratings from FQ to FQ+1 trials (*t*_19_ = 3.17; *P* < .0001) but no such difference for ON ratings (*t*_19_ = 2.07; *P* > .1).

### III-Evolution of ON vs. OFF state

To investigate the dynamics of MW with time on task (see ***Figure 1E***) we compared MW levels between FQ+1 and all FQ+N trials. The trial duration allows a definition from ON to OFF with a temporal resolution of ∼ 2 sec. ON and OFF probabilities show a negative (*r* = -.64; q = .049) and positive (*r* = .81; q = .014) effect with time on task, respectively. Pairwise comparisons between FQ+1 and all FQ+N trials (FQ+2,…,FQ+25) show statistical significance with q < .01 (see ***Figure 1F***). The difference in OFF ratings between FQ+1 and FQ+2 indicates a drift from ON to OFF within at least 2 seconds.

### IV-Performance as a function of ON vs. OFF state

We examined whether performance changed between trials with reported labels (the labels provided by the participants) and predicted labels (predicted by the SVM). Performance differed between brain states (*F*_2,17_ = 10.5, *P* < .0001; see ***Figure 2A***) but not between label types (*F*_1,17_ = .021, *P* = .88). The interaction effect between brain state and label type was not significant (*F*_1,17_ = .0053, *P* = .99). Neither OFF (*t*_19_ = 1, *P* = .32; 58.1 (SD = 24.6) vs. 58.0 (24.6) %) or MID (*t*_19_ = .41, *P* = .68; 74.7 (16.5) vs 74.1 (16.0) %) nor ON (*t*_19_ = 1.08, *P* = .23; 77.6 (19.0) vs. 76.7 (18.3) %) performance differed between label types. Both the reported and predicted labels performance differed between ON vs. OFF (reported: *t*_19_ = 3.32, *P* = .0036; predicted: *t*_19_ = 3.19, *P* = .0048) and ON vs. MID (reported: *t*_19_ = 2.72, *P* = .013; predicted: *t*_19_ = 2.9, *P* = .0091). We then compared sub- and suprathreshold performance. Importantly, the perception thresholds were comparable across brain states (OFF: 5.78°, MID: 6.57 °, ON: 5.82°; see ***Figure 2B***). We averaged the performance values separately for all brain states for both the subthreshold (1.5 - 4.5°) and suprathreshold (7.5 - 15°) tilt angles. We found a trend for a main effect of brain state (*F*_2,17_ = 2.84, *P* = .06; see ***Figure 2C***) and a significant main effect for threshold level (*F*_2,17_ = 30.28, *P* < .0001) but this effect was due to a difference between OFF vs MID (*t*_19_ = 3.35, q = .0061) and OFF vs ON (*t*_19_ = 3.08, q = .0067) performance only for sub-threshold stimuli. But importantly, stimulus discrimination was not different between the ON and OFF states (all *t*s < 1; all qs>.1). Mean reaction times differed between brain states (*F*_2,17_ = 3.93; *P* = .02; see ***Table 1*** and ***Figure 2D***) but not between threshold levels (*F*_2,17_ = 2.41; *P* = .12). The variance of reaction times ς RT differed between brain states (*F*_2,17_ = 9.52; *P* < .001; see ***Table 1*** and ***Figure 2E***) but not between threshold levels (*F*_2,17_ = 2.31; *P* = .13).

**Table 1:**
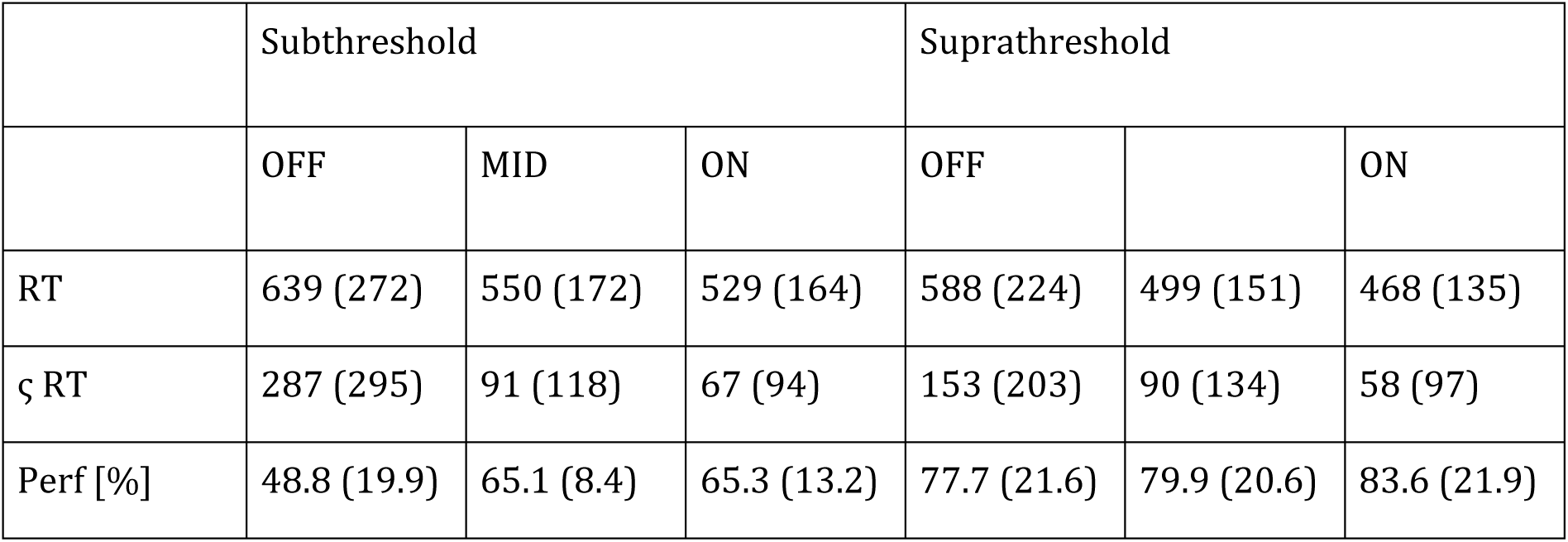
mean performance measures (standard deviation in parentheses): reaction time (RT), the variance of reaction times (ς RT), and discrimination performance in % for OFF and ON trials, separately for sub- and suprathreshold stimulation.

**Figure 2.**
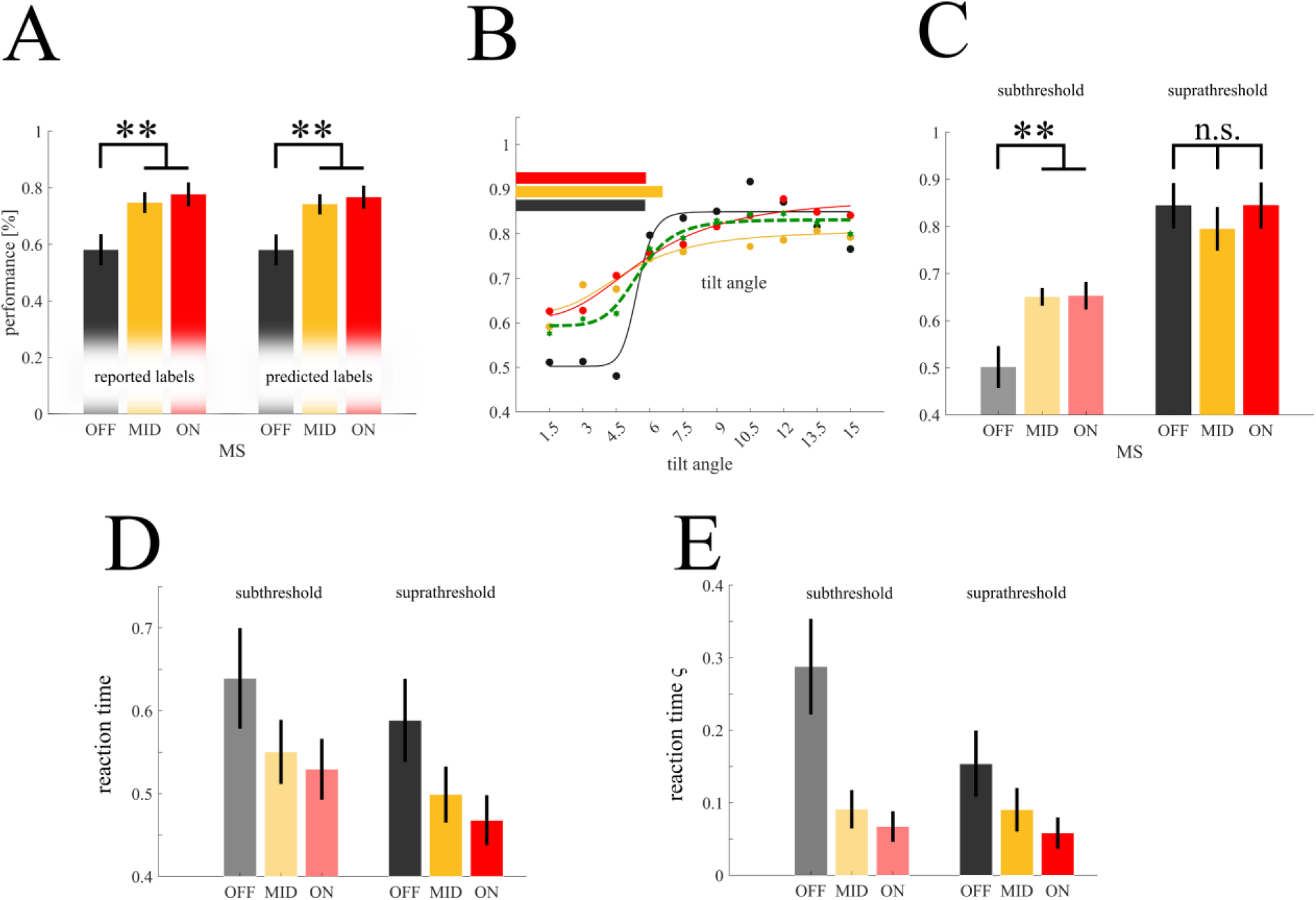
Behavioral results: ***A*** Target discrimination accuracy grouped by reported (left) and predicted (right) labels shows a performance decrease in the OFF condition. ***B*** shows performance curves for the three states (OFF: black, MID: yellow, ON: red) with similar perceptual thresholds across conditions (horizontal bars). ***C*** We compared performance between brain states and threshold levels (sub- vs suprathreshold) and found that OFF performance differed from MID and ON performance in the subthreshold but not in suprathreshold stimuli. ***D*** and ***E*** show that reaction times and the variance of reaction times ς, respectively, are increased in OFF trials compared to MID and ON trials both in the subthreshold (left) and suprathreshold (right) trials.

### V-Stimulus Response

The BHA in MEG sensors covering the occipital cortex (see ***Figure 3A***) showed z>3 between 116 and 208 msec after stimulus onset (see ***Figure 3B***).

**Figure 3.**
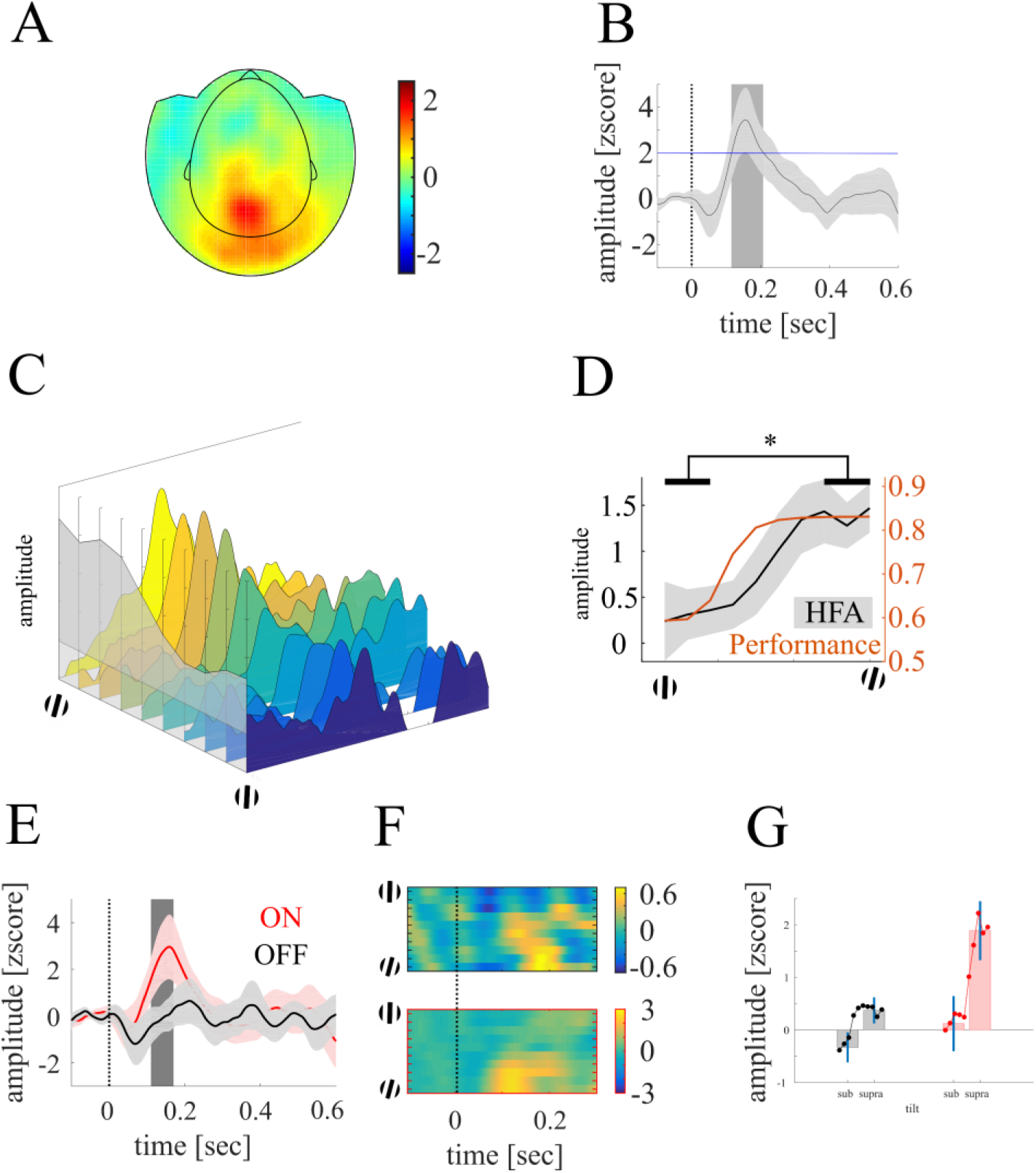
Depiction of the BHA response to visual stimulation. **A** shows the topographic distribution of BHA averaged across the temporal interval shown in B. **B** shows the time course of BHA recorded at occipital MEG sensors with a z-value >2 (stimulus-responsive channels). **C** shows the BHA in stimulus-responsive channels as a function of the orientation angle of the target. For example, the yellow and dark blue areas show the BHA to the largest and smallest orientation angle, respectively. The grey area in the foreground shows the maximum BHA values to each of the different orientation angles. High and low orientation angles lead to high BHA and low BHA, respectively, in logistic form. **D** shows BHA averaged across the interval of significant modulation (see gray shaded area in B). The solid line and shaded area show mean and standard error, respectively, across all subjects. The orange line shows the performance curve averaged over all trials. **E** shows the BHA for both ON (red line) and OFF (black line) trials. The dashed line marks the stimulus onset. The gray shaded area in the background shows the temporal interval of significant difference between ON and OFF. The solid line and shaded area show mean and standard error, respectively, across all subjects. **F** shows the BHA in response to the different orientation angles for both OFF (upper) and ON (lower panel) trials. In both groups, large orientation angles correspond to high BHA in logistic form, although overall BHA shows a stronger modulation in ON than OFF trials. **G** shows the BHA in OFF (black) and ON (red) trials for both subthreshold and suprathreshold stimuli.

### VI-BHA variation as a function of target orientation

The BHA was larger in trials with a suprathreshold compared to a subthreshold orientation angle (*t*_19_ = 2.48; *P* = .022; see ***Figure 3C&D***) indicating that BHA encodes the target orientation. Next, we examined whether BHA also varied with MW.

### VII-BHA variation with brain state

In stimulus responsive channels, we found stronger BHA modulation in ON compared to OFF trials between 112 to 168 ms (*t*_crit_ = 2.16; *t*_max_ = 2.48; q = .038, see ***Figure 3E***). In the last step we tested for BHA variation as a function of target orientation both in ON and OFF trials. (see ***Figure 3F***). In both brain states, we observed a larger BHA in suprathreshold orientation angles compared to subthreshold orientation angles (OFF: *t*_19_ = 1.89; *P* = .04; ON: *t*_19_ = 1.87; *P* = .04; see ***Figure 3G***).

In experiment 2, using simultaneous electroencephalographic-magnetoencephalographic (EEG-MEG) recordings in 23 participants, we examined the stimulus locked temporal kinetics of BHA modulation and whether the BHA varied with brain states. We compared the BHA to the initial visual feedforward sweep to V1, indexed by the EEG-C1 component. The C1 is also linked to an increase in multi-unit activity in visual sensory areas ^17–21^. If BHA is a measure only for feedforward visual input, a prediction would be that BHA and C1 overlap in latency. Participants were presented with a visual detection paradigm with a target arrow pointing to the left or right side, accompanied by high contrast checkerboard wedges to elicit a robust C1 (see ***Figure 4A***). Participants had to respond to the orientation of the arrow via button press. As in experiment 1, thought probes were presented pseudorandomly in 20% of the trials and participants rated their level of attention to the trial immediately preceding the probe. We first examined whether participant’s behavioral performance varied as a function of brain states. We further compared the latencies of EEG-C1 and MEG-BHA and finally analyzed whether the components varied with brain states (see ***Results VIII – XII***).

**Figure 4.**
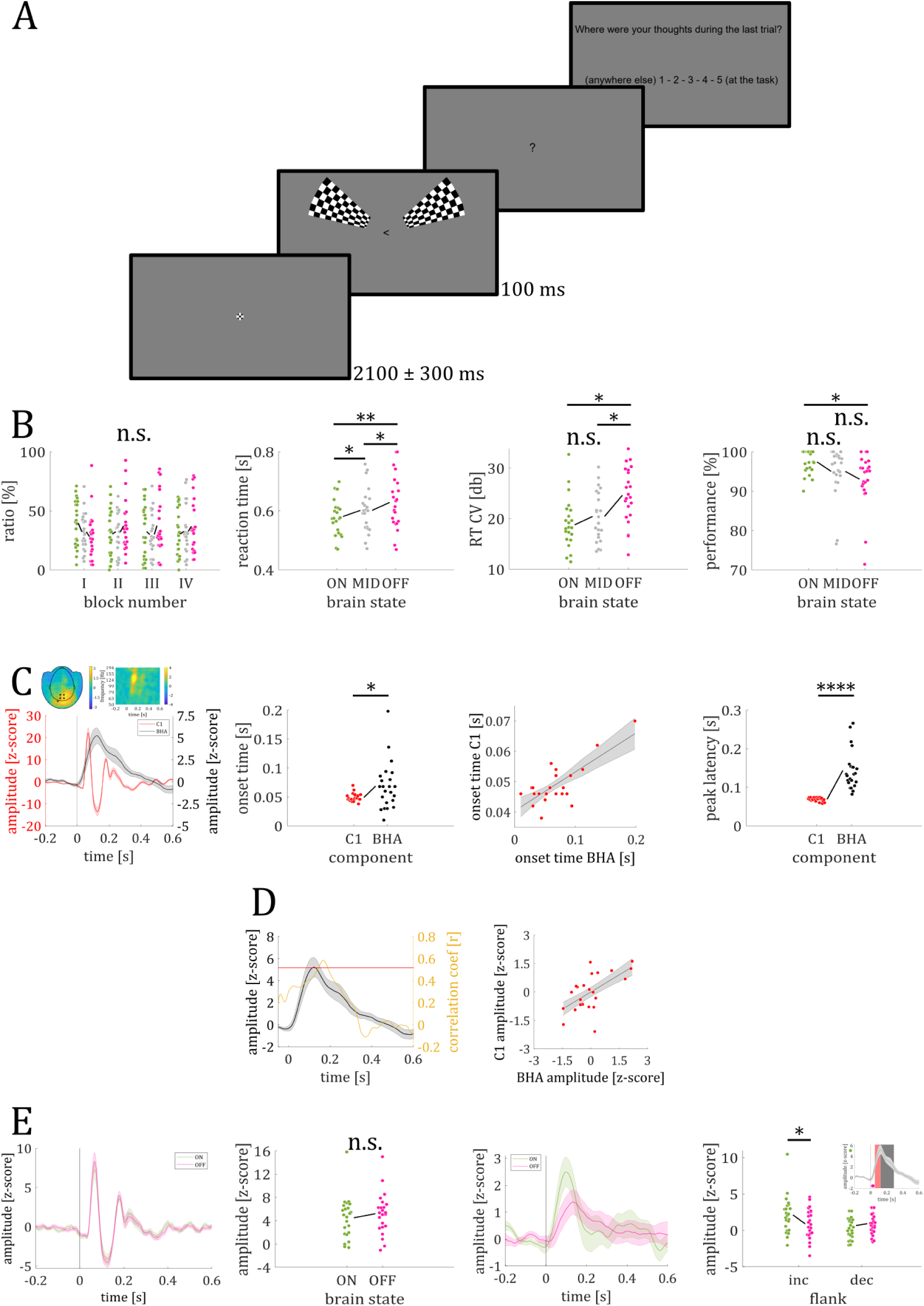
Paradigm and results of experiment 2. ***A***: visual stimulation with high contrast checker-boards accompanying the visual discrimination task. In 25% of the trials a thought probe was presented upon which participants had to their current level of focus on the task. ***B***: Likelihood of brain states did not vary across the blocks (left). Reaction times changed with brain states with slowest and more variable response during OFF trials. Subjects made most errors during OFF trials (right). ***C***: Time course of grand average C1 and BHA response (left). Topographic plot and time-frequency representation show spatial and frequency-specificity of the BHA response. Both onset and peak of C1 were earlier than onset and peak of BHA. The scatter plot shows the strong correlation of BHA and C1 onset times. ***D***: shows prediction of BHA magnitude by C1 peak response. Black line shows BHA (black) and yellow line time-resolved correlation coefficient. Red line shows 99% confidence interval for correlation coefficient value. BHA correlated significantly with peak C1 response in the time range between 140 ms and 192 ms (left). Scatter plot shows maximal correlation between BHA and peak C1 response. ***E:*** C1 does not show variation with brain state (ON – green; OFF – magenta) Individual mean C1 amplitude for ON and OFF trials (second from left). BHA shows variation with brain states in the time interval of the increasing but not decreasing flank (see inset).

### VIII – Behavioral Results

#### Likelihood of brain states

Thought probes were presented on average every 16.51 s (*SD* = 1.35s; range: 14.57 s – 20.12 s). The 4 x 3 ANOVA with the factors block (I, II, III, IV) and brain state (ON, MID, OFF) showed no significant main effect of block (*F*_3, 262_ = 0.01; *P* = .998), brain state (*F*_2, 262_ = 0.58; *P* = .56; see ***Figure 4B***) nor significant interaction effect (*F*_6, 262_ = 1.01; *P* = .42), indicating that the frequency of the different brain states did not differ across the experiment (see ***Figure 4B***).

#### Reaction times

Participants’ reaction times differed between brain states (*F*_2, 44_ = 12.83; *P* < .0001; see ***Figure 4B***). Post-hoc *t*-tests revealed longer reaction times during OFF than ON (*M_OFF_* = 628 ms; *M_ON_* = 581 ms; *t*_22_ = 3.91; *P* < .001) and MID trials (*M_MID_* = 602 ms; *t*_22_ = 2.94; *P* < .01), as well as longer reaction times during MID than ON trials (*t*_22_ = 3.50; *P* < .01). Similarly, we found a significant main effect (*F*_2, 44_ = 10.54; *P* < .001; see ***Figure 4B***) when we compared RT CV values between brain states (ON, MID, OFF). RT CV values for OFF trials were higher than for ON (*M*_OFF_ = 24.64; *M*_ON_ = 18.79; *t*_22_ = 3.55; *P* < .01) and MID trials (*M*_MID_ = 20.44; *t*_22_ = 3.60; *P* < .01), but no difference was found between ON and MID trials (*t*_22_ = 1.57; *P* = .13).

#### Target discrimination performance

Participants’ performance differed between brain states (F_2, 44_ = 6.27; *P* < .01; see ***Figure 4B***). Post-hoc *t* tests revealed worse performance in OFF compared to ON trials (*M_OFF_* = 93.2 %; *M_ON_* = 97.5 %; *t_22_* = 3.09; *P* < .01), but no performance difference between OFF and MID trials (*M_MID_* = 95.0%; *t*_22_ = 1.72; *P* = .10), nor between ON and MID trials (*t*_22_ = 2.13; *P* = .05). Note, that overall performance was very high across brain states.

### IX – C1 Response

The C1 component peaked at the posterior EEG electrodes PO3 and PO4. Since time-resolved t-tests showed no significant differences between the two EEG channels (PO3_max_ = 3.35 µV; PO4_max_ = 3.04 µV; all *P* > .13), we averaged the C1 across both sensors in the subsequent analysis. Checkerboards in the UVF and LVF elicited a negative and positive C1 response at posterior electrodes, respectively. Since absolute C1 amplitude values did not differ between UVF and LVF (*M_UVF_* = 4.14 µV; *M_LVF_* = 5.32 µV; *t*_22_ = 1.95; *P* = .06), we converted the negatively trending C1 component for UVF stimulation into a positive one by multiplying the EEG traces by -1. We then averaged the resulting time series across UVF and LVF trials to obtain a single C1 response. Comparing the collapsed ERP time course with a surrogate distribution revealed a significant C1 time interval from 46 to 88 ms (zC1_crit_ = 1.48; zC1_max_ = 22.20 at 68 ms; all *P*-values < .00001; see ***Figure 4C***). Pupil dilation did not differ between brain states in the C1 time interval (mean zPD_ON_ = 2.38; mean zPD_OFF_ = 0.72; *t*_13_ = 1.18; *P* = .26).

### X – Broad Band High-frequency Activity

Five MEG sensors showed a significant BHA amplitude modulation between 50 ms and 288 ms after stimulus onset compared to a surrogate distribution (z_crit_ = 3.00; BHA_max_ = 5.22 at 122 ms; all *P*-values < .041; see ***Figure 4C***).

### XI – C1 and BHA latency comparison

We found both a significant earlier onset of C1 than BHA (C1_onset_ = 49 ms; BHA_onset_ = 69 ms; *t*_22_ = 2.62; *P* = .02; see ***Figure 4C***) and onset times were strongly correlated (*r* = .76; *P* < .0001; see ***Figure 4C***) indicating that this effect can be seen on a single-subject level. Furthermore, the C1 peaked significantly earlier than the BHA (C1_peak_ = 68ms; BHA_peak_ = 144 ms; *t*_22_ = 7.07; *P* < .00001; see ***Figure 4C***) but peak times were not correlated (*P* = .43). The peak C1 amplitude at 68 ms correlated with BHA amplitude between 140 ms and 192 ms (*r*_crit_ = .50, *r*_max_ = .58 at 166 ms; *P* = .002; see ***Figure 4D***).

### XII – Amplitude Modulation with brain state

C1 amplitude did not differ between brain states (*t*_22_ = 1.02; *P* = .32; see ***Figure 4E***). However, the increasing flank of the BHA (50 – 122 ms) showed a significant difference between brain states with higher BHA amplitude in ON compared to OFF trials (*t*_22_ = 1.99; *P* = .03; see ***Figure 4E***). No such difference was found for the decreasing flank (122 – 288 ms; *t*_22_ = 0.57; *P* = .71; see ***Figure 4E***).

## Discussion

We examined how brain states changes during wakefulness affect human perception and early visual response. We compared the effects of ON state (focused wakefulness) with the OFF state (inattention/mind-wandering). In the first experiment we used a classifier that relied on theta activity in MEG sensors covering visual cortex to distinguish between brain states. We found that humans shift rapidly from an ON to an OFF state, with transitions occurring within a few seconds. These rapid state transitions did not impact target detection to suprathreshold stimuli but had slower and more variable reaction times during the OFF state. Importantly, we discovered that broadband high-frequency activity (BHA) recorded by MEG sensors in the occipital cortex correlated with target orientation, providing a marker for perceptual performance. Although the BHA was reduced during the OFF state, the extraction of sensory information supporting target detection using BHA amplitude remained intact. In the second experiment we investigated whether broadband high-frequency activity (BHA) reflects input to visual cortex and whether the BHA amplitude modulation is primarily influenced by bottom-up information or by internal brain states. We used the EEG C1 response as a reference measure for initial sensory input to the visual cortex. The BHA peaked later and was more sustained than the C1 and was modulated by internal brain states.

In contrast to the C1 component, we observed a decrease in the amplitude of the BHA during periods of mind wandering. Why did we observe a modulation of BHA by the internal brain state given that primate research suggests that the BHA response reflects input to visual cortex ^22^? Using intracranial recordings of V1 in macaque monkeys, Leszczyński et al. ^10^ investigated BHA response and its relationship to multi-unit activity. The authors found the supragranular BHA was the largest contributor to the recorded surface BHA, which was interpreted that the surface recorded BHA includes significant cortical feedback representations ^10^. The different response characteristics of EEG-C1 and MEG-BHA we observed support this interpretation. The BHA peaks later than the C1 component supporting the view that the BHA contains representations of cortical feedback processes which are reduced during mind wandering.

This temporal lag between the C1 and BHA is in line with studies on feedback projections to V1. Previous EEG studies exploring the flow of visual information along the dorsal and ventral pathways observed activity in dorsolateral frontal cortical regions approximately 30 ms after the onset of the C1 component ^23^. Based on this rapid system-wide activation, it was suggested that early feedback processes could be initiated, leading to early feedback projections to V1 within the time range of N1 and P1, roughly around 100-200 ms ^23^. These results align with intracranial recordings conducted in macaque monkeys, which revealed context-dependent alterations in V1 activity that did not correspond to the traditional properties of V1 cell receptive fields. Notably, these contextual modifications were observed as early as 30 ms after the initial activation of V1 ^24,25^. To summarize, these findings suggest that corticocortical feedback projections to V1 can commence as early as 30 ms after the initial activation of V1.

Individual trial OFF-state classification using MEG was consistent in the theta range, aligning with previous EEG studies suggesting stronger distinctions between brain states in lower frequencies ^3,26,27^. A few studies have trained classifiers to delineate brain state changes ^28–30^. These studies can be categorized into two approaches: individual-level predictions or predictions across multiple participants. Jin et al. employed convolutional neural networks (CNNs) to train EEG classifiers for monitoring the OFF state using EEG data collected during a visual search task and the Sustained Attention to Response Task (SART). However, the authors acknowledged the challenge of generalizability across participants. Dong et al. ^29^ examined ERPs in an oddball paradigm and attempted to generalize the findings to multiple participants. They reported that individual-level decoding accuracy was higher, than across participants, supporting Jin et al.’s findings. In our study, we adopted an approach similar to Mittner et al. ^28^, who used fMRI and operationalized the OFF-state using introspective thought-probes. We and Mittner et al. ^28^ trained a support-vector machine to classify each individual trial as on-task or off-task based on subjects’ responses to the probes. Consequently, each trial was assigned a probability indicating the likelihood of the participant being on-task or off-task. Mittner et al. achieved a decoding accuracy rate approaching 80% in a two-class problem using fMRI. We achieved an accuracy of 63% for 3 classes using MEG. Collectively, the extant research indicates that brain states can be reliably predicted using fMRI, EEG, and MEG activity.

Our study indicates rapid shifts from an ON to an OFF state among human subjects, often occurring within two seconds and found reliable MEG decoding at the single-trial level. Focus queries are sporadically and randomly administered to minimize the risk of reducing mind wandering ^31^. However, this poses a challenge for EEG investigations since a limited number of trials results in suboptimal signal-to-noise ratio. As a result, brain activity in previous studies is averaged over several preceding trials, assuming implicitly that participants were in the same brain state. These results were obtained using either a fixed number of preceding trials ^29,30^ or a specific time duration, such as several seconds^3^. Another approach involves selecting different numbers of trials preceding OFF and ON ratings ^26^. In this study, mind wandering trials were defined as those in which participants reported mind wandering, along with the two preceding trials, while the remainder of trials were categorized as on-task trials. However, this approach is susceptible to mixing different brain states, as demonstrated by the rapid shifts between brain states we observed. Here we show that participants tend to spend less time in the ON state than commonly assumed. The practice of averaging over previous trials introduces the mixing of brain states and the rapid variation in brain states we observed needs to be taken into account during state decoding.

One could argue that brain state decoding of single trials is an indirect measure since we relied on participants’ report in only 20% of the trials. Our behavioral findings support the validity of our data-driven estimation of brain states. First, the predicted ON and OFF labels are in accord with the distribution of reported ON and OFF labels. Second, the behavioral markers of target detection associated with brain states are consistent between reported and predicted trial classes. The overall performance across all sub- and suprathreshold stimuli also shows a comparable pattern for reported and predicted labels. In both trial sets, the performance was worse in OFF trials compared to ON trials, which is notable considering factors like performance and reaction times were not included in the classification. Our findings using brain activity decoding show that behavioral differences between ON and OFF states can be reliably determined.

Across both experiments the differences between ON and OFF states are most evident in reaction times and reaction time variability in line with previous studies showing variations in motor responses ^32,33^. For example, Leszczynski et al. ^34^ conducted a SART task requiring participants to continuously monitor a stream of stimuli presented at the center of a computer screen. Subjects pressed the space bar whenever a non-target item appeared but withheld the button-press response when a target item was displayed. During mind wandering the authors report an increase in both average reaction time and reaction time variability similar to our findings. The authors suggest that mind wandering has a detrimental effect on performance, as evidenced by changes in reaction times. In contrast, we confirmed that reaction time is increased during MW, but threshold detection performance was intact in line with previous studies^35^. In conclusion, in both experiments we could show that during mind wandering, we are able to perceive stimuli effectively, although with longer and more variable response times.

We found high-frequency activity in MEG sensors covering the visual cortex resembling the BHA response observed in previous studies using intracranial recordings. Notably, in V1 (primary visual cortex), a strong onset response modulation is observed within a time window of less than 200 milliseconds, with a peak around 100 milliseconds ^16,36–38^. Since MEG primarily captures tangential sources in the cortex and high frequencies have less spatial spread, we assume that the source of our BHA signal is located mainly in V1. We found that the BHA exhibits two patterns varying both with the degree of target grating orientation and the behavioral state. Invasive studies on animals and non-invasive recordings in humans using magnetoencephalography (MEG) consistently demonstrate that high-contrast grating stimuli elicits gamma band activity, typically ranging from 30 to 100 Hz, in the local field potential (LFP) recordings of primary visual cortex (V1) ^39,40^ and that the amplitude of this activity varies with orientation ^41–44^. In humans, the frequency of the visual gamma response also correlates with orientation discrimination thresholds across individuals ^45^. The definition of the gamma band varies with studies, so higher frequencies (>80Hz) are analyzed along with low gamma without differentiating between the two.

However, recent studies have shown that brain responses in this frequency range exhibit different response characteristics ^38^. The BHA band also responds to more complex stimuli, while lower frequency gamma activity responds mainly to gratings. Our study expands the existing previous results by demonstrating that BHA can be reliably measured using MEG and that the amplitude of BHA indicates the strength of the orientation of the target gratings.

Previous EEG studies have shown reduced sensory responses, primarily in the low-frequency range (<30 Hz), during mind wandering (MW) ^6,7,46^, suggesting decreased sensory input. Visual BHA is responsive to the physical properties of visual stimuli and provides another metric to study sensory processing across brain states ^38^. Studies in animals have primarily focused on brain state changes during locomotion, although inattentive states have also been described ^47^, contrasting with alert wakefulness. These three states - locomotion, alert wakefulness, and inattention - differ primarily in arousal levels, with locomotion inducing the highest arousal and the inattentive state exhibiting the lowest. The behavioral state plays a role in how the brain responds to visual stimuli, beginning with the thalamus ^47^ and early processing stages depend on corticothalamic feedback, which is reduced during inattention ^48^. Conversely, higher arousal levels, such as those experienced during locomotion, lead to increased visual responses ^49^. To date, no studies have demonstrated how the inattentive state influences the response in early visual cortex, particularly in terms of BHA changes, neither in animals nor humans. Here, we demonstrated that BHA is reduced during the OFF state. However, reduction in BHA amplitude does not reduce target orientation discrimination, in line with previous studies ^16^. These authors examined the neurophysiology of sustained visual perception and found, despite a significant reduction in BHA, the ability to decode categories was maintained. We extend these results by demonstrating changes in the BHA onset response to basic visual stimuli but preserved target orientation discrimination.

Our results demonstrate that human visual perception is not static but exhibits substantial drifts on a second-to-second macroscale. Previous research has shown microscale sub-second fluctuations in perception at a 4 Hz rate in both macaques and humans^50^ with reaction times varying at this cycle rate. Here we show similar perceptual fluctuations by revealing macroscale drifts in visual perception over seconds.

In conclusion, we found that human subjects undergo rapid transitions from an ON to an OFF state, occurring within a few seconds. These transitions do not affect the ability to differentiate visible targets but result in slower and more variable reaction times during the OFF state. We provide insight into the relationship between BHA and behavioral states, showing that BHA amplitude is influenced by both target orientation and the brain state. While previous studies have focused on the effects of arousal levels on visual responses, our findings contribute to understanding how internal attention states influence the response during the OFF state in early visual processing stages. These observations highlight the dynamic nature of brain states during wakefulness and their impact on perception and early visual response.

## Methods

### Experiment 1

Experiment 1 examined the connection between BHA and variations in target orientation and whether this association remained consistent across different brain states. Initially, we performed a single-trial classification to identify the frequency band that most accurately reflected the individual ratio of ON and OFF states to test transition times between ON and OFF.

### Participants

20 subjects (9 female, range: 19-39 years, M: 29.11, SD: 6.28) participated after providing written informed consent. All participants reported normal or corrected-to-normal vision, and none reported any history of neurological or psychiatric disease. All recordings took place at the Otto von Guericke University of Magdeburg and were approved by the local ethics committee (“Ethical Committee of the Otto-von-Guericke University Magdeburg”). Participants received monetary compensation.

### Paradigm

Participants were presented with a stimulus array of red, green, and blue grating patterns each consisting of three colored stripes displayed as if viewed through a circular aperture (***Figure 1A***). The grey stripes matched the background color. Either green or red gratings served as target and blue gratings always served as distractor items. Stimulus arrays consisted of 18 gratings arranged in two blocks of 9 gratings, one in the left lower visual field and the other in the right lower visual field. Participants were instructed to keep fixation on the fixation cross located at 1.9° visual angle (va) above the stimulus array. The size of each grating was 1.15° va and distance between single gratings (edge-to-edge) was 0.69° va. The left and right block of gratings each had a size of 4.83° by 4.83° va, the horizontal distance between both blocks (inner edges) amounted to 5.15° va. Diagonal distance between the fixation cross and the center of the nearest upper grating was 2.81° va. Target gratings were tilted left or right in ten steps of 1.5°, with the smallest tilt being 1.5° and the maximal tilt being 15° relative to the vertical axis. Orientation and tilt angle of the non-target and distractor gratings varied randomly across trials.

### Procedure

At the beginning of each block (n=12), participants were instructed to attend either to the red or green grating and report via button press which side it was tilted to (left: index finger, right: middle finger of the right hand). Target color assignment alternated blockwise. In blocks with the red grating as target the green grating served as non-target to be ignored and vice versa. Targets could appear at each of the 18 locations. The location of the non-target was constrained to the mirrored location in the opposite grating block to maintain equal distances to the fixation cross for both target and non-target gratings. Each trial started with a fixation period of 1250 ms (±250ms) before the stimulus array was presented for 100 ms. Participants were asked to respond as fast and accurately as possible. In total, one trial lasted 1.91 sec (SD = .09 sec). This means that we can resolve the evolution of ON to OFF for a temporal resolution over two second intervals. The experiment started with a training block of twenty trials to familiarize participants with the procedure. After twenty consecutive trials, a seven second pause allowed participants to blink and rest their eyes. Each block consisted of 100 trials.

### Experience Sampling

Thought probes (focus queries; *FQ*) were presented pseudorandomly (20% of all trials) and participants rated their level of attention to the trial immediately preceding the probe on a five-point scale from 1 (”thoughts were anywhere else” – OFF) to 5 (”thoughts were totally at the task” – ON). Responses to questions were given using all five fingers of the left hand (thumb: 5, index finger: 4, middle finger: 3, ring finger: 2, little finger: 1). The probes were presented following orientation discrimination, with the restriction that two probes were separated by a minimum of one intermediate trial (this minimal distance of one intermediate trial between probes occurred for 7% of the focus queries). The probes were initiated by an auditory stimulus (500 Hz, ca. 85 dB for 200 msec). Note that an increased number of thought probes (>20%) would decrease the time between single probes reducing the time for mindwandering.

### Decoding of Brain States

#### Preprocessing

We extracted 27 frequency bands with exponentially increasing center frequencies from 3 to 180 Hz to determine the frequency band that best predicted brain states (see ***Figure 1B***). Each band had a bandwidth equal to 30% of its corresponding center frequency. In the preprocessing step, we employed the MNE Python package ^51^ to perform band-pass filtering and extract 1-second epochs of each trial. These epochs spanned from -.3 to .7 seconds relative to stimulus onset. We concatenated the time series of all MEG sensors within an epoch into one vector representing the feature space. We then normalized across the epochs to a normal distribution *N* (0,1) for two reasons. First, normalization reduces the dominance of high variance features over low variance features. Second, a support vector machine using a radial basis function as kernel requires data with zero mean and variance of the same order.

#### Comparison of Machine Learning Algorithms

To select the machine learning algorithm with best decoding performance, we compared (see ***Table 2***) Linear Discriminant Analysis (LDA) ^52^, Logistic Regression (LR) ^53^, and Support Vector Machines (SVM) ^54^. We performed classification on single-participant and multi-participant levels for each of the three methods, see ***Table 2*** and ***Table 3***, respectively. All data sets with reported labels were split into training and evaluation sets with an 80:20% ratio. Five-fold cross-validation was used for classification accuracy.

**Table 2:** Summary of model accuracy of single participant analysis.

|  | Accuracy in % |  |  |
| --- | --- | --- | --- |
| Participant | LR | LDA | SVM |
| 1 | 48.48 | 43.94 | 50.00 |
| 2 | 65.28 | 68.06 | 68.06 |
| 3 | 37.50 | 48.61 | 50.00 |
| 4 | 45.07 | 46.48 | 47.89 |
| 5 | 37.50 | 50.00 | 50.00 |
| 6 | 63.38 | 59.15 | 59.15 |
| 7 | 83.10 | 84.51 | 83.10 |
| 8 | 50.70 | 49.30 | 50.70 |
| 9 | 87.50 | 87.50 | 87.50 |
| 10 | 51.39 | 55.56 | 56.94 |
| 11 | 49.30 | 57.75 | 59.15 |
| 12 | 93.06 | 95.83 | 93.06 |
| 13 | 52.78 | 38.89 | 51.39 |
| 14 | 48.61 | 52.78 | 56.94 |
| 15 | 58.33 | 55.56 | 45.83 |
| 16 | 54.93 | 56.34 | 57.75 |
| 17 | 94.44 | 94.44 | 94.44 |
| 18 | 47.89 | 57.75 | 54.93 |
| 19 | 66.67 | 72.22 | 70.83 |
| 20 | 66.20 | 70.42 | 73.24 |

**Table 3:** Summary of model accuracy of multi-participant analysis.

|  | Number of train data |  |  | Number of test data |  |  | Accuracy in % |  |  |
| --- | --- | --- | --- | --- | --- | --- | --- | --- | --- |
| Number of Participants | OFF | MID | ON | OFF | MID | ON | LR | LDA | SVM |
| 20 | 469 | 1104 | 1725 | 184 | 479 | 751 | 51.06 | 52.47 | 60.54 |

**Table 4:**
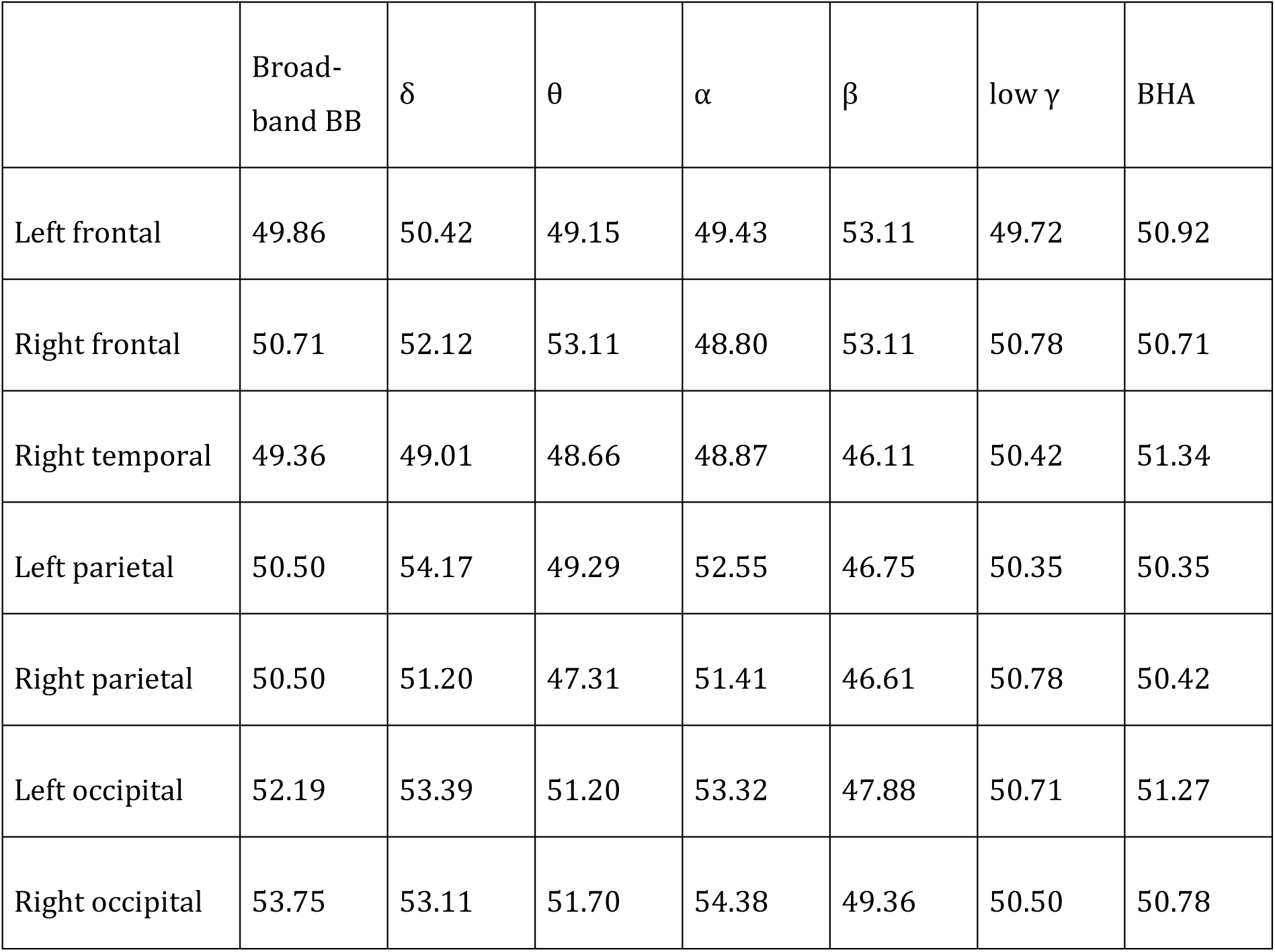
Summary of SVM accuracy separately for canonical frequency bands and regions of interest.

The SVM was as the best-performing method among the three. Note that the three approaches were fine-tuned with specific adjustments. In the case of LDA, the SVD solver was chosen. LR underwent regularization with L2, and the penalty term C was set at 0.01. The following section is dedicated to detailed description of the selected SVM method.

#### Support Vector Machine

SVM is well suited for MEG recordings since it can perform with a limited training data in high-dimensional spatio-temporal space. The epoched data, transformed into a feature vector, is then subjected to SVM analysis. We trained on all 20 subjects and 5 categories in a multi-participant analysis. The classifier is implemented in Scikit-Learn ^55^, where SVM is based on LIBSVM ^56^. The multi-class support is handled according to a one-vs-one scheme.

We fine-tuned hyperparameters in the SVM model by performing a grid search with cross-validation to enhance classification performance and handle imbalanced classes. Unlike optimizing on a participant level, which we deemed impractical, we adopted a multi-participant approach. This process involved assessing a predefined list of parameter values and dividing the data into folds, ensuring that each fold had a comparable distribution of classes to the entire dataset. We tested the parameter permutation of regularization parameter *C* ∈ {0.01, 0.05, 0.1, 1, 10} and kernel ∈ {linear, RBF, sigmoid}. For the multi-participant prediction, the resulted best performing choice was *C equal* 1 using the RBF kernel. For the RBF kernel, free parameter γ was selected as 1/(*N_(features)_* ∗ *σ_data_*).

### Statistical Analysis Experiment 1

We focused on the relationship between BHA and changes in target orientation and whether this relationship was constant over brain states. We first conducted a single trial classification to determine the frequency band that best represented the individual ratio of ON and OFF states (*I-Representation of individual ON vs. OFF ratio*). Following that, we investigated the effectiveness of thought probes in interrupting OFF-state performance and bringing participants back to an ON state after a focus query (*II-Effectiveness of thought probes*). Subsequently, we explored the temporal changes in the state following the focus query (*III-Evolution of ON vs. OFF state*). We compared participants’ performance in different brain states (*IV-Performance as a function of ON vs. OFF state*) and examined the relationship between BHA and both target orientation and brain state. We identified MEG channels displaying a response to stimuli (V*-Stimulus response*). Finally, we assessed whether the BHA stimulus response varied depending on the level of target orientation (*VI-BHA variation as a function of target orientation*) and brain state (*VII-BHA variation with brain state*).

### I-Representation of individual ON vs. OFF ratio

Decoding accuracy provides information about the separability of brain states. Another criterion is the degree of prediction of interindividual differences in levels of ON vs OFF state. Previous studies report significant individual differences between participants in the ratio of ON and OFF ratings (R_FQ_) assessed with focus queries (FQ). Since FQs were presented at random time points, the R_FQ_ for these trials serves as a proxy for the individual overall ratio. We then investigated how well the R_FQ_ from predicted labels of the individual frequency bands reflected the individual ratios. First, we determined the individual likelihood distribution of labels 1 to 5 reported at the FQ. We then determined the same frequency distribution of labels 1 to 5 using the predicted labels R_PL_ for each frequency band. Finally, we determined the deviation Δ_R_ of R_PL_ from R_FQ_ for each frequency band using the root mean square.

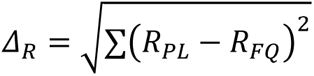

Frequency bands with minimum deviation of predicted ON/OFF ratings best represented individual MW level.

### II-Effectiveness of thought probes

An implicit assumption is that MW in the OFF state is interrupted when FQ are presented, allowing participants to report on their prior thought content. However, the effectiveness of thought probes is unclear ^57^. A direct way to test the effectiveness of the FQ method is to compare the MW ratios reported with focused queries (FQ trials) with the predicted MW labels in the trials immediately afterwards (FQ+1 trials). Under the condition that the FQ effectively interrupts MW, we assumed that the frequency of OFF predictions (made by the classifier) in FQ+1 trials was smaller than OFF ratings (reported by the subjects) in FQ trials. In contrast, we assumed a different pattern with smaller or no changes in ON ratings between FQ and FQ+1. To test this, we calculated the likelihood of OFF ratings (rating 1 – 2) and ON ratings (rating 4 – 5) for each participant separately for FQ and FQ+1 trials. Note that the likelihood distribution of FQ trials represents the distribution of reported labels while the likelihood distribution in FQ+1 trials is calculated on predicted labels. We compared the likelihood values with a 2-way ANOVA with the factors trial type (FQ vs. FQ+1) and brain state (OFF vs. ON) across participants. The observed F values were compared against a surrogate distribution. The surrogate distribution was constructed in the following way. In 1000 runs, we calculate the likelihood of OFF ratings (rating 1 – 2) and ON ratings (rating 4 – 5) for each participant as explained above. For the FQ+1 trials we calculated across as many randomly chosen trials as there were FQ+1 trials. This leads to 1000 surrogate F values both for the main and interaction effect. We separately assigned a *P*-value to the F-value of the main effect within the surrogate distribution of the main effect and a *P*-value to the F-value of the interaction effect within the surrogate distribution of the interaction effect.

### III-Evolution of ON vs. OFF state

Previous state dependent research required participants to indicate whether their mind wandered or not across a period spanning several trials. However, it is not clear how stable the ON state is and at which temporal scale we return to the OFF state after ON task performance. The machine learning approach and the resulting predicted labels enabled us to estimate a time course of the transition from an ON task to OFF task state. To test the evolution of the MW state, we compared the probability of being in the ON-state across consecutive trials after the actual FQ. In each trial after the FQ (FQ+N), we determined the probability for ON and OFF states as described above. We estimated the ON/OFF ratio for all trials ranging from FQ+1 through FQ+25, since the maximum number of intervening trials between two consecutive thought probes was 25 (see ***Figure 1***). Since individual trials lasted ∼2 sec we were able to test the evolution of MW across 50 sec with a temporal resolution of 2 sec.

We then tested for the effects of ON and OFF ratings across FQ+N trials. The probability values for ON and OFF ratings were averaged across subjects resulting in a time series for ON and OFF ratings. In a first step, we tested for a variation of the averaged ratings over the FQ+N trials using Pearson’s correlation coefficient *r*. We then tested whether this correlation is different from zero by comparing observed *r*-values against a surrogate distribution. In 1000 runs, we calculated the likelihood of OFF and ON ratings for each participant and predicted labels were randomly assigned to single trials. In each iteration we estimated Pearsońs *r* leading to 1000 surrogate *r* values. We assigned a *P*-value to the observed *r*-values within the distribution of surrogate *r*-values. Both a significant negative *r* of ON ratings and a significant positive *r* of OFF ratings over time indicate an increase in the likelihood of OFF state. The correlation provides a trend over time but does not define how the probability distributions between successive trials differ across participants. In a second step we tested for differences between successive trials and hence steps of 2 sec. To increase the statistical power, we formed a common measure using ON and OFF probabilities separately for each FQ+N. Since ON probability is disproportionally higher than OFF (see ^58^), the time course of ON and OFF ratings was standardized (z-score) separately for each subject. Since OFF ratings have a significant increase across time and ON ratings have a significant decrease (see Results), OFF ratings were multiplied by -1 and then averaged with ON ratings for each subject, separately for each FQ+N trial. We then compared the MW in FQ+1 trials with all FQ+N trials across subjects with a t-test. The comparison of observed difference values with the surrogate distribution results in a *P*-value for each pairwise comparison. To control for multiple testing, we corrected the *P*-values by applying the false discovery rate (FDR) ^59^ across all pairwise comparisons. The adjusted *P*-values are labeled q. *T*-value with a q < .05 (corrected *P*-value) were classified as showing a significant difference.

### IV-Performance as a function of ON vs. OFF state

Previous MW studies on reading and the Sustained Attention to Response Task (SART) report performance decline during MW ^60,61^ but examination of the influence of the OFF state on visual perception is lacking. Further, focus queries are sporadically and randomly administered to minimize the risk of reducing the likelihood of mind wandering ^31^ which would reduce the number of trials for each target orientation. We investigated how the threshold changes using predicted trials. We first tested whether predicted labels reflect the differences in performance between brain states as seen using reported labels. Note, that for the performance we used all ratings (OFF: rating 1 and 2; MID: rating 3; ON: rating 4 and 5). Using a two-way ANOVA with the factors brain state (OFF, MID, ON) and label type (reported vs. predicted) we compared performance measured as percent correct responses averaged across tilt angles, across subjects. We did not find differences between predicted and reported labels (see ***Results***) and merged both label types (reported and predicted) in the next steps investigating how brain states fluctuations affect perception (target discrimination) and reaction times across the tilt angles.

The predicted labels allow us to determine a mean threshold across subjects for ON, MID and OFF. We fit a logistic function to the data separately in each condition

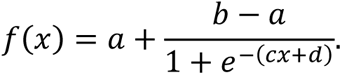

The perceptual threshold we defined as the x value (target orientation angle) corresponding to 75% correct discrimination. We then tested whether differences between brain states were more likely due to performance differences below or above the perceptual threshold. We compared performance values in a two-way ANOVA with the factors brain state (OFF, MID, ON) and threshold level (sub- vs supra-threshold). Post-hoc we compared performance between OFF, MID, and ON both for sub- and suprathreshold across subjects using *t*-test. The comparison of observed *t*-values with the surrogate distribution results in a *P*-value for each pairwise comparison. The surrogate distribution was calculated in the following way. In 1000 runs we randomly assigned labels (OFF, MID, ON) to the individual performance values and calculated *t*-values as explained above. To control for multiple testing, we corrected the *P*-values by applying the false discovery rate (FDR) across all pairwise comparisons. The adjusted *P*-values are labeled q. *T*-value with a q < .05 (corrected *P*-value) were classified as showing a significant difference. Mean reaction times (RTs) and variance of reaction times were grouped for the OFF and ON states and threshold level (sub- vs. suprathreshold) and averaged within subjects. The RTs where then compared using a two-way ANOVA with the factors brain state (OFF, ON) and threshold level (sub- vs. suprathreshold) across participants.

### V-Stimulus Response

We identified stimulus-responsive channels showing a significant BHA modulation (z>3) following the onset of the visual search display for at least 100 msec using the following steps. We z-scored the BHA values as described below and determined in the baseline interval of each trial and each channel both the mean amplitude 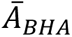 and the standard deviation ς_BHA_. We then subtracted 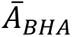 from the BHA and divided the result by ς_BHA_ at each time point in each trial and channel. Channels with BHA modulation (averaged across trials) z>3 following the onset of the visual presentation across at least 100 msec were considered stimulus responsive.

### VI-BHA variation as a function of target orientation

We first tested whether the BHA in stimulus responsive channels represents target orientation. In our study, this was target orientation angle. We grouped trials by the absolute angle of the target and averaged the BHA time series across trials and stimulus-responsive MEG sensors. We then compared BHA in response to subthreshold (1.5°–4.5°) and suprathreshold targets (7.5°– 15°; see also *Performance as a function of MW*) in the temporal interval of significant stimulus response. The resulting values were compared across subjects using a *t*-test.

### VII-BHA variation with brain state

In the next step, we investigated whether BHA varied with ON and OFF state. We grouped trials by predicted labels into OFF trials (labels 1 – 2) and ON (labels 4 – 5) and averaged the baseline corrected time series across trials and stimulus-responsive MEG sensors in each subject. We then compared the BHA time series across participants between the ON and OFF condition using a *t*-test. We compared each *t*-value against a surrogate distribution constructed under the null assumption of no difference between ON and OFF. This surrogate distribution was constructed by randomly reassigning the labels (ON vs. OFF) to the single trial averages in 1000 permutations providing a1000 surrogate t-value time series. We assigned a *P*-value to each *t*-value within the surrogate distributions. The *P*-values for *t-*value were corrected for multiple testing by applying the FDR. F and r with q < .05 were classified as significant.

We then examined whether BHA independent of absolute strength also represented target orientation. For this we compared the BHA response to sub- and suprathreshold stimuli between ON and OFF trials across subjects. We grouped trials by the absolute angle of the target item for both the ON and OFF condition. We then averaged the BHA time series across trials and stimulus-responsive MEG sensors and compared BHA in response to subthreshold in the temporal interval of significant difference between ON and OFF. The resulting values were compared across subjects using a directed *t*-test.

### Experiment 2

In experiment 2 we tested the stimulus locked temporal kinetics of BHA modulation and whether the BHA is modulated differently during ON and OFF states. To address this, we compared the behavior of BHA to the initial feedforward sweep to V1. Specifically, we focused on the C1 event-related potential component in the electroencephalography (EEG), which reflects the initial feedforward volley to visual cortex. The C1 component, recorded at occipital EEG electrodes, peaks prior to 100 ms after stimulus onset, serving as a marker for initial afferent activity in the primary visual cortex ^62^. The C1 is also linked to an increase in multi-unit activity in visual sensory areas ^17–21^. If BHA is a measure only for feedforward visual input, a prediction would be that BHA and C1 overlap in latency.

### Participants

23 subjects (17 female, range: 18-35 years, M: 26.58, SD: 4.95) participated after providing written informed consent. All participants reported normal or corrected-to-normal vision, and none reported any history of neurological or psychiatric disease. All recordings took place at the Otto von Guericke University of Magdeburg and were approved by the local ethics committee (“Ethical Committee of the Otto-von-Guericke University Magdeburg”). Participants received monetary compensation.

### Stimuli

We employed a visual detection paradigm with high contrast checkerboard wedges ^63^ to elicit a robust C1 (see ***Figure 4A***). Black arrows presented in the center of the screen functioned as targets and were accompanied by the checkerboards, all stimuli were presented against a grey background. The targets had a size of 1.5 cm x 0.7 cm, covering 0.54 ° of visual angle (va), while the checkerboards had a size of 12 cm x 10 cm, covering 8.02 ° of va. The target was accompanied by bilateral high-contrast black-and-white checkerboard wedges either in the upper (UVF) or lower (LVF) visual field. Each wedge was part of a ring-like checkerboard with a spatial frequency of 5 cycles per degree with each wedge covering 3.125 % of the area of the checkerboard ^63^. The checkerboards in the UVF and LVF were located at polar angles of 25 ° above and 45 ° below the meridian to stimulate the lower and upper banks of the calcarine fissure ^62,64^.

### Procedure

Each trial started with the presentation of a fixation cross for 2.1 s (± 300 ms) the subjects were instructed to look at. After this period the target arrow appeared at the fixation location for 100 ms pointing to either the left or right side.

Participants identified the direction of the arrow and responded as fast as possible via button press with the right index (for left) or middle finger (for right). The experiment consisted of four blocks with 272 trials in each block, resulting in a total number of 1088 trials. The paradigm was counterbalanced, so that in 50% of the trials a checkerboard appeared in the UVF and in 50% of the trials the checkerboard appeared in the LVF in a pseudorandomized order. The arrow pointed to the right in half of the trials and to the left in the other half of trials. The experiment was programmed and controlled with Psychtoolbox 3 ^65^ running on Matlab (R2018b).

### Experience sampling

In 20% of the trials in experiment 2 the stimulus array was followed by a thought probe in which we asked participants to rate their attentional focus in the trial immediately preceding the thought probe on a 5-point Likert-scale from 1 (“thoughts were anywhere else”—OFF) to 5 (“thoughts were totally at the task”—ON). Responses to these probes were given with all 5 fingers of the left hand (little finger: 1, ring finger: 2, middle finger: 3, index finger: 4, thumb: 5). Thought probes were presented in a pseudo-random order with the restriction that two consecutive probes had to be separated by at least one trial. We limited the number of thought probe trials to enhance the likelihood of Mindwandering. A recent study indicates that eight to ten thought probes are sufficient to gain reliable and valid information about the Mindwandering rates ^66^. To increase statistical power, we grouped the five mind wandering ratings in three groups of brain state (OFF: 1 & 2; MID: 3; ON: 4 & 5). We incorporated only the trials that directly preceded the thought probes in our examinations.

### Statistical Analysis Experiment 2

We first compared the behavioral measures for different brain states in experiment 1 (*VIII-Behavioral Performance*). This included both reaction times and accuracy, defined as the percentage of correct responses, and then assessed the distribution of mind wandering. We analyzed the EEG C1 component (*IX-C1 Component*). Next, we analyzed the MEG BHA component (*X-Broad Band High Frequency Activity*). In the following step, we compared the latencies of the EEG-C1 and MEG-BHA and analyzed whether C1 peak response predicted participants’ BHA amplitude (*XI – Latency differences between C1 and BHA*). Finally, we analyzed whether the EEG-C1 and MEG-BHA varied with brain states (*XII-Amplitude Modulation with experimental condition*). We compared each statistical parameter against a surrogate distribution. To determine the time interval of grand average response of C1 and BHA, we compared the observed data against a surrogate distribution. In 1000 iterations, we constructed the surrogate distribution of time series by circularly shifting time series of participants between -1 and 2 s separately. We then compared the original data with this surrogate distribution by calculating the cumulative distribution function, providing a *P*-value for each time point indicating whether there was a significant amplitude modulation over baseline. To determine time intervals of statistically significant differences between brain states, we compared each statistical parameter against a surrogate distribution, that was calculated by randomly yoking the labels of the trials and repeating the analysis in 1000 iterations. Thus, all *P*-values reflect the significance relative to the surrogate distribution.

### VIII – Behavioral Performance

#### Reaction times

Trials with extreme reaction times < 100 ms or > 2000 ms were marked as outliers and excluded. Reaction times were then grouped for the three brain states, averaged across subjects and compared using a one-way ANOVA with the factor brain state (ON, MID, OFF). In addition, we computed reaction time coefficient of variation (RT CV) for every participant for the three brain states. RT CV is a measure of RT variability while controlling for RT speed and is computed by dividing the RT’s standard deviation by the mean RT and multiplicated with the factor 100 (Epstein et al., 2011). The resulting RT CV values were compared using a one-way ANOVA with the factor brain state.

#### Target discrimination performance

Performance was determined for each subject. The resulting performance values were compared between brain states (ON, MID, OFF) using a one-way ANOVA.

#### Likelihood of brain states

We tested whether the ratio of brain state reports changed over the experiment to rule out the possibility that changes in cortical dynamics are a result of a change across the experiment and not due to brain state fluctuations throughout the experiment. We averaged participant’s brain state ratings for each block and compared the ratio of brain state ratings across the blocks using a 4 x 3 ANOVA with the factors block (I, II, III, IV) and brain state (ON, MID, OFF).

### IX – C1 Response

We identified the EEG-C1 by labelling trials as a function of position of checkerboards presentation (LVF vs. UVF). Stimulation in the LVF generates a positive going C1 and stimulation in the UVF generates a negative going C1. C1 component amplitude did not differ between UVF and LVF (see ***Results IX – C1 Response***). Hence, responses to UVF stimulation were multiplied by -1 to revers polarity and then averaged with LVF stimulation in each channel and subject. To identify the C1 window (amplitude modulation over baseline), we compared the C1 activity in posterior EEG channels PO7/8 against a surrogate distribution. *P*-values were then corrected for multiple comparisons, using the False Discovery Rate (FDR) method ^59^.

### X – Broadband High Frequency Activity

We then analyzed the BHA for each trial and magnetometer. We band-pass filtered the time series in the broadband high frequency range (80–150 Hz) and extracted the analytic amplitude Af (t) of this band by Hilbert-transforming the filtered time series. In the following analysis BHA refers to this Hilbert transform. We smoothed the time series so that the amplitude at a given time point t was always the mean activity of 20 ms around this time point. The data were then baseline corrected subtracting the mean activity of the time interval 200 ms preceding stimulus onset from every data point for each trial and each channel. Afterwards we identified channels showing a significant BHA amplitude modulation following stimulus presentation. We z-scored BHA amplitude values for each channel by calculating the mean and standard deviation of the BHA in the baseline period compared to the post-stimulus 300 msec window. First, we subtracted the mean from the BHA at each time point and divided the BHA at each time point by the standard deviation. Channels with z>3 (corresponding to *P* = .0027) in the interval ranging from 0 to 300 ms were labeled as stimulus responsive. We then averaged the BHA data over the stimulus responsive MEG channels and determined the time window when the BHA showed a significant modulation by comparing each observed time point with the surrogate distribution.

### XI – C1 and BHA latency comparison

To analyze whether the C1 peak response predicted participants’ BHA amplitude, we first z-scored C1 and BHA amplitude values. We calculated participants’ individual time points of C1 and BHA response onset defined as the time the C1 and BHA amplitude first showed a z>3. This results in an onset time point for each subject for both the C1 and BHA. We then compared C1 and BHA response onset times using a *t*-test and calculated Pearson’s correlation coefficient of C1 and BHA onset times. We then identified participants’ individual time points of C1 and BHA peak response and the amplitude value at this time point. Peak responses were compared using a *t*-test and a Pearson correlation coefficient of C1 and BHA peak times. We tested whether C1 amplitude magnitude predicts BHA amplitude and calculated Pearson correlation coefficients between C1 amplitude at peak time point (∼at 68ms) and BHA amplitude values at each time point separately. We compared these correlation coefficients with a surrogate distribution. In 1.000 iterations, we took the C1 and BHA values of the subjects at the time of the highest observed correlation, randomly reassigned the C1 and BHA values to the subjects and then calculated the Pearson correlation coefficient. We then calculated the 99% Confidence Interval for the correlation coefficient resulting in a critical coefficient value. Correlation coefficients exceeding this critical value were considered as showing a significant correlation between C1 peak response and BHA amplitude.

### XII – Amplitude Modulation with experimental conditions

We analyzed differences between brain states in C1 and BHA separately using a *t*-test. We z-scored the C1 and BHA time series by calculating the mean and standard deviation of the C1 in the baseline period. We subtracted the mean from the C1 at each time point and divided the C1 at each time point by the standard deviation. The same was done for the BHA. Since the BHA response is known to have a fast onset and a slowly decreasing flank, we compared mean amplitudes of the increasing BHA flank and the decreasing BHA flank separately.

## Funding

This study was partially supported by the German Research Foundation - DFG grant DFG SFB-1436, TPA03.

## Author contributions

S.D. and P.S. conceived and designed the experiment. S.D. and P.S. collected the MEG data. P.S., T. K., P. M., C.R., S. S., and S.D. analyzed the data, P. S., T. K., P. M., C.R., S. S., R.T.K. and S.D. interpreted the data. R.T. K., P. S. P. M., C.R., S. S., and S.D. wrote the manuscript.

## Competing Interest Statement

No competing interests

## Classification

Biological Sciences – Neuroscience

## Notes

### Competing Interest Statement

The authors have declared no competing interest.

